# pH - responsive, reversible A-motif based DNA hydrogels: synthesis and biosensing applications

**DOI:** 10.1101/2022.12.06.519399

**Authors:** Vinod Morya, Ashish Kumar Shukla, Chinmay Ghoroi, Dhiraj Bhatia

## Abstract

Functional DNA hydrogels using various motifs and functional groups require perfect sequence designing to avoid cross-bonding interference with self or other structural sequences. The present work reports an A-motif functional DNA hydrogel that does not require any sequence design. A-motif DNA is a non-canonical parallel DNA duplex structure comprises homopolymeric deoxyadenosines (poly-dA) strands that undergo conformational changes from single strands at neutral pH to a parallel duplex DNA helix at acidic pH. Despite many advantages over other DNA motifs like no sequence, design is required and no cross-bonding interference with other structural sequences, A-motif has not been explored much. We successfully synthesized DNA hydrogel utilizing A-motif as a reversible handle to polymerize DNA three-way junction (3WJ). The composed A-motif hydrogel was first characterized by EMSA, & DLS, which shows the formation of higher-order structures. Further, we utilized imaging techniques like atomic force microscopy (AFM) and scanning electron microscope (SEM) validating its hydrogel like highly branched morphology. pH-induced conformational transformation from monomers to gel is quick and reversible, and was analysed for multiple acid-base cycles. The sol-to-gel transitions and gelation properties is further examined using rheological studies. The use of A-motif hydrogel in the visual detection of pathogenic target nucleic acid sequence is demonstrated for the first time using the capillary assay. Moreover, the pH-induced hydrogel formation is observed *in-situ* as a layer over the mammalian cells. The proposed A-motif DNA scaffold has enormous potential in designing stimuli-responsive nanostructures that can be utilized for many biological applications.

## 1. Introduction

DNA-based hydrogels, where DNA is used as the structural material or cross-linker are now known for two decades. DNA Hydrogels are highly branched hydrophilic polymers with a vast scope of functionality. They have shown a plethora of scope in various fields like biosensing^1^, controlled drug release^2^, tissue engineering^3^, etc.^4,5^ The properties of the hydrogel depend upon the polymer used and the degree of cross-linking. Stimuli-responsive, reversible cross-linker makes the hydrogel ‘smart’, which changes its properties at certain predefined environmental conditions or responses to small molecules or ionic cues^6^. There are many examples of stimuli-responsive DNA structures that follow Watson-crick base paring as well as non-Watson-crick base paring^7^. To make the hydrogels functional, different motifs and functional groups have been used, including aptamers^8^, i-motifs^9^, G-quadruplexs^10^, antibodies^11^, small peptides^12^ etc. These motifs and functional groups work as actuators upon specific stimuli. Cytosine rich DNA forms i-motif at acidic pH, which is quadruplex in nature^13^. Although i-motifs have been explored for many applications in the field of DNA nanotechnology^14^, but they require sequence designing. These motifs need an interference check with other DNA structures. Moreover, during designing the sequence, as they can hybridize with self or the other sequences and form unwanted DNA secondary structures. In contrast, A-motif DNA hydrogel is a potential alternative to the pH responsive i-motifs.

A-motif is a parallel duplex DNA helix consisting of homopolymeric deoxyadenosines (poly-dA)^17^. The adenosine in the poly-dA single strands undergoes protonation and pairs at low pH by A^+^H-H^+^A, the conformational changes are reversible^17^. There is slight structural difference compare to B-DNA, the distance between the strands in the A-motif helix is higher than the B-DNA i.e., 2.2 nm^17^. Saha et al., employed A-motif to make a rigid 1D DNA architecture and demonstrated the reversible properties^18^. Considering the simple and non-interfering sequence of poly-dA, first time we explored its properties for making a stimuli-responsive, reversible DNA hydrogel. We introduced the A-motif forming overhangs on pre-defined B-DNA three-way junction (3WJ) monomers to make a reversible DNA hydrogel, responsive to pH conditions.

In the present work, we have synthesised and characterized the A-motif cross-linked DNA hydrogel. 3WJ with poly-dA overhangs on each arm designed to polymerise and form a 3D matrix. These strands self-assemble to form 3WJ via Watson-Crick complementary base-paring. (**Figure 1a**). The hybridization of poly-dA at acidic pH crosslink the 3WJ monomers leads to formation of hydrogel like architecture. These 3WJ remain as monomer at neutral pH and polymerizes to form hydrogel only at acidic pH (<5). The formation of higher order structures by pH change was characterized by electro mobility shift assay (EMSA) and dynamic light scattering (DLS) spectroscopy. The atomic force microscopy (AFM) and scanning electron microscopy (SEM) were used to analyse the morphology. The viscoelastic properties of the formed A-motif hydrogel were studied on a rheometer.

**Figure 1.**
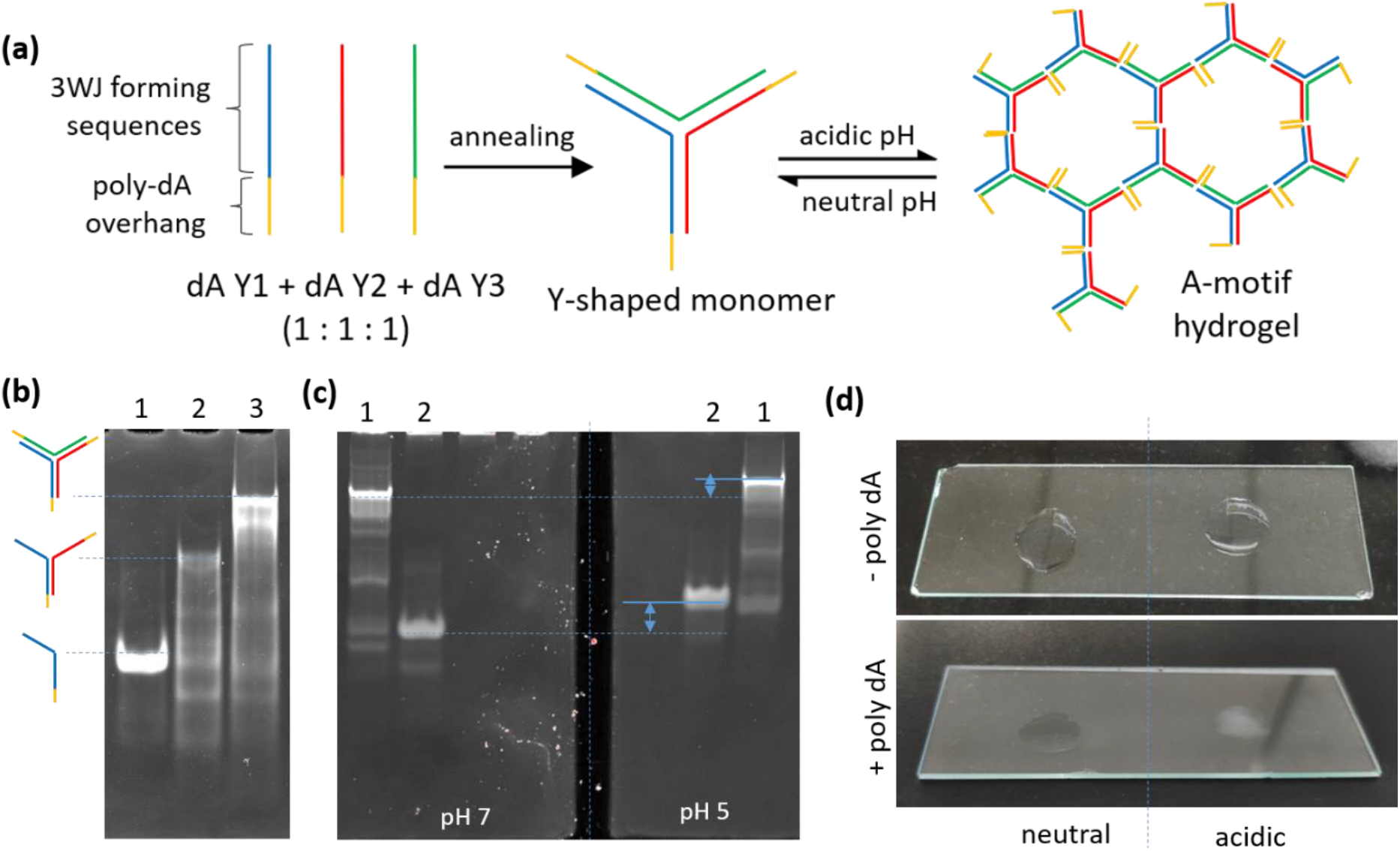
Design, synthesis, and conformation of A-motif hydrogels. (a) Schematic of self-annealing process of 3WJ with poly-dA overhang followed by pH induced hydrogel formation. (b) Gel electrophoresis showing the retardation in the mobility upon formation of 3WJ using 10% native-PAGE; Lane1: dAY1, 2: dAY1 + dAY2, 3: dAY1+ dAY2 + dAY3. (c) Gel electrophoresis showing the retardation in the mobility at acidic pH (1x BRB, pH5) in comparison to neutral (1x BRB, pH7) using 8% native-PAGE; Lane 1= dAY1+ dAY2 + dAY3, 2=dAY1. (d) Visuals of 3WJ with and without poly-dA overhang at different pH, 3WJ with poly-dA overhang become little opaque in acidic condition.

In addition to the preparation of pH responsive hydrogel we also tested in-situ hydrogel formation on the surface of mammalian cells to show its consistency and feasibility in the biological system. We further utilized the physical properties of the A-motif DNA hydrogel i.e., sol-to-gel formation for a biosensing application. The sol-to-gel formation leads to the increase in water holding capacity, which significantly decrease the capillary action-induced fluid rise in a capillary tube. Lastly, this phenomenon availed to make a visual detection system to detect target nucleic acid.

## 2. Results

### 2.1 Design and synthesis

The sequences for monomer 3WJ (Y-shaped structure) were adapted and modified from previously reported hydrogels^19,20^ (**Supplementary information, Table S1**). As, the minimum required length to form an A-motif is six nucleotide^17^, we decided to keep it twelve to make sure the formation of rigid A-motif. Two random nucleotides were added in between the poly-dA tail and the 3WJ forming sequence to give flexibility to the overhangs. The 3WJ monomers with poly-dA overhang were formed via self-assembly of three sequences named as dA-Y1, dA-Y2 and dA-Y3 and confirmed by EMSA (**Figure 1b**). To form the hydrogel, 0.2 mM HCl was added to the monomer’s solution to make the final pH 5. At acidic condition, the poly-dA overhangs undergo conformational changes and form A-motif by A^+^H-H^+^A, which polymerises the 3WJ monomers in a hydrogel. Decreasing pH for the single strand with poly-dA overhang results in formation of a homodimer; which shows the formation of A-motif at acidic condition. The decrease in mobility can be seen at the acidic condition in comparison to neutral pH condition (**Figure 1c**). Moreover, the formation of hydrogel can be seen visually as an increment in the optical density at acidic pH^21^ (**Figure 1d**).

### 2.2 Characterization

#### 2.2.1. Formation and reversibility of hydrogel

The structural changes in terms of hydrodynamic diameter at different pH can be observed by DLS. The peak around 10 nm at neutral pH showing the smaller particle size i.e., monomers 3WJ, while in acidic pH the larger particles (>100 nm) can be seen which indicates the formations of higher order structures (**Figure 2a**). The change in hydrodynamic sizes were monitored on multiple acid-base cycle and found the reversible sol-to-gel transition up to 3 cycles (**Figure S1**).

**Figure 2.**
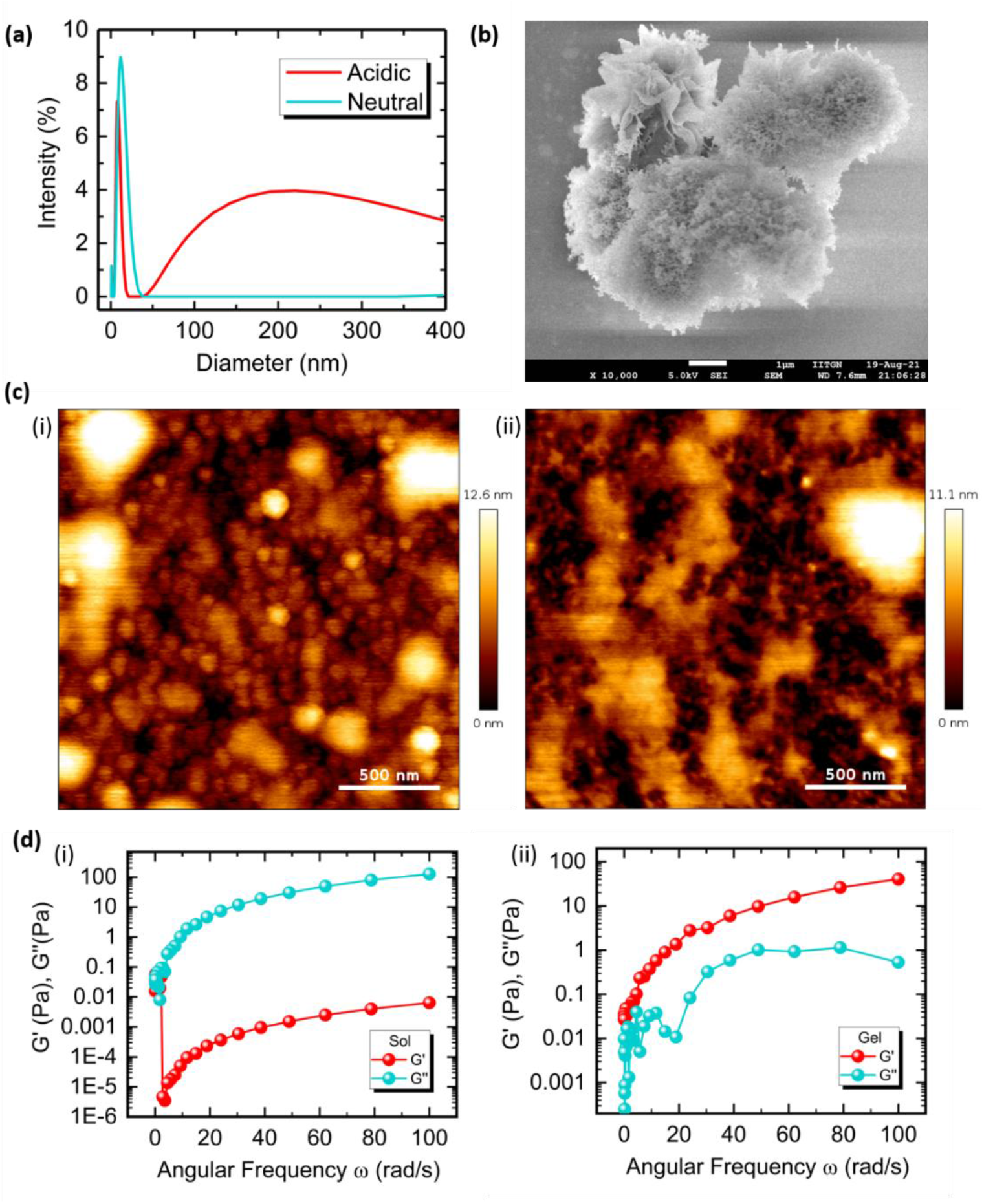
Characterization of As-motif hydrogel at different pH conditions. (a) The hydrodynamic diameter characterized by DLS showing the monomers (<10 nm) at pH 7 and formation of higher order structure (>100 nm) at pH 5. (b) FE-SEM image of lyophilized A-motif hydrogel (scale bar = 1 μm). (c) AFM images of (i) A-motif hydrogel forming monomers at pH 7, and (ii) A-motif hydrogel at pH 5. (d) Comparison of storage modulus (G’) and loss modulus (G”) as a function of angular frequency (ω) at (i) pH 7 and (ii) pH 5; representing typical behaviour of sol and gel respectively.

#### 2.2.2. Morphology characterization at different pH conditions

The morphological characterization of the hydrogel was performed on FE-SEM and AFM. The FE-SEM images of the freeze-dried hydrogel samples showed a highly branched globular structure of micro-hydrogel (**Figure 2b**), similar to previously reported DNA hydrogels^21^. The AFM images of individual particles at neutral pH dictates the monomeric form and a highly branched network with fissures at acidic pH confirming the porous structure of hydrogel (**Figure 2c**).

#### 2.2.3. Rheological studies

Rheological studies have been performed to know the visco-elastic properties of the pH-responsive hydrogel. The rheological experiments were carried out using a freshly prepared monomer solution for A-motif hydrogel. A hydrogel refers to a gel, when elastic response (G’, storage modulus) should be greater than viscous response (G″, loss modulus)^22^. The elastic (G′) and viscous response (G″) response of the A-motif hydrogel were examined as a function of shear strain at 25°C, in the angular frequency range 0.1-100 rad s^−1^, as per previously reported method^21^. At neutral pH the G’ is lesser than the G” (**Figure 2d**); while at acidic pH, the G′ was approximately an order of magnitude higher than the G″ (**Figure 2e**). This indicates the elastic nature of the A-motif hydrogel increase in acidic condition. Additionally, we observed that the G′ and G″ have not crossed each other, which indicates that the hydrogel is stable and rigid^23^. As a result, A-motif hydrogel responds better to gelation at acidic pH since its elasticity is dominant over its viscosity. These observed features confirm the assembly of 3WJ with poly-dA tail in to hydrogel with change in pH from neutral to acidic.

### 2.3 *In-situ* formation of hydrogel on the surface of mammalian cells

The formation of hydrogel was not limited to solution phase in a tube, we have tried to mask the cell membranes with A-motif hydrogel formation *in-situ*. To achieve this, one of the three stands (dA Y1-Cy5) in the 3WJ monomer was tagged with Cyanine 5 (Cy5) fluorophore. HeLa cells. The cells were allowed to grow for 24 hrs post seeding, then 5 μM as working concentration of 3WJ-poly-dA-Cy5 in PBS buffer with pH 7 were added after removing the media. Cells were incubated for 10 minutes; to let them sit on the outer cell membrane, after that the buffer was replaced by pH 5 PBS buffer. The fixed cells at both the pH were imaged and found that at pH 7 the fluorescence was very discrete and of low intensity, while at pH 5 the fluorescence intensity increased significantly and mesh like structure also can been seen around the cells (**Figure 3a**). The quantified intensity shows more than a double increase in the fluorescence (**Figure 3b**), which signifies the in-situ formation of A-motif hydrogel at acidic pH.

**Figure 3.**
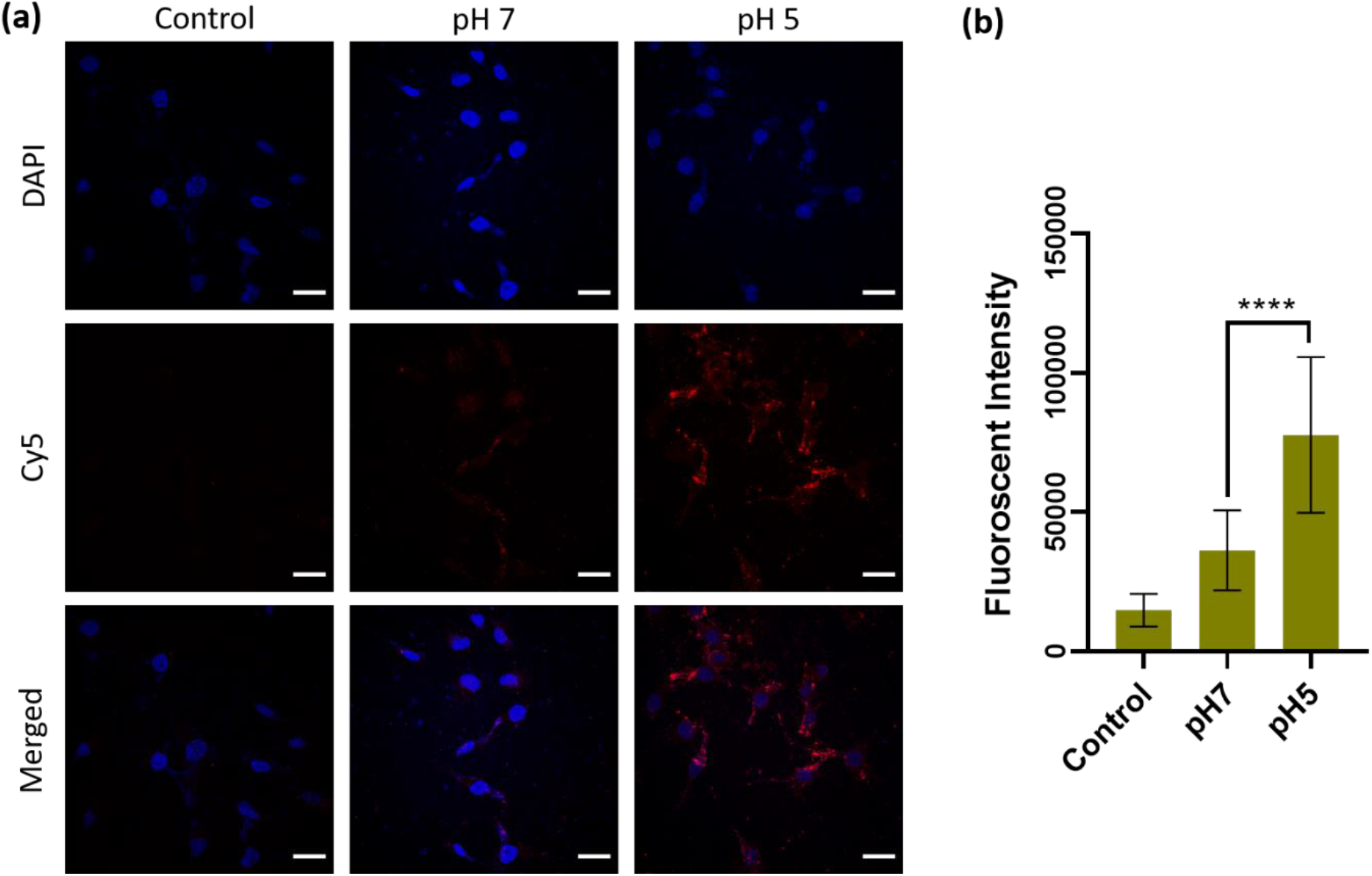
*In-situ* formation of A-motif hydrogel on the surface of mammalian cells. (a) Confocal images of HeLa cells introduced with A-motif forming monomers at different pH. The red channel (Cy5) showing discrete fluorescence at neutral pH and increase in the fluorescence intensity at acidic pH (scale bar= 10 μm). (b) Quantification of fluorescence intensity at different pH. Error bars indicate standard deviation; n = 30 cells per condition (ordinary one-way ANOVA, P value <0.0001****).

### 2.4 Visual detection of target DNA by capillary assay

#### 2.4.1 Design of CHA driven A-motif hydrogel for capillary assay

The visual detection of a specific target DNA were carried out by catalytic hairpin assembly (CHA) driven 3WJ formation and pH induced capillary action. The three strands of 3WJ designed in such a way that they form hairpin-loop individually with poly-dA overhang at 5’ end and one of the strands in 3WJ complimentary to the sequence of target DNA. The three sequences i.e. A1, A2, and A3 annealed individually by heating up to 95°C and rapidly cooled to 20°C to make stable hairpin-loop. Strand A1 was complementary to the target DNA (TD), and in presence of TD, the hairpin-loop disrupt and make a duplex with TD by toehold-mediated strand displacement^24^ (**Scheme 1a**).

**Scheme 1.**
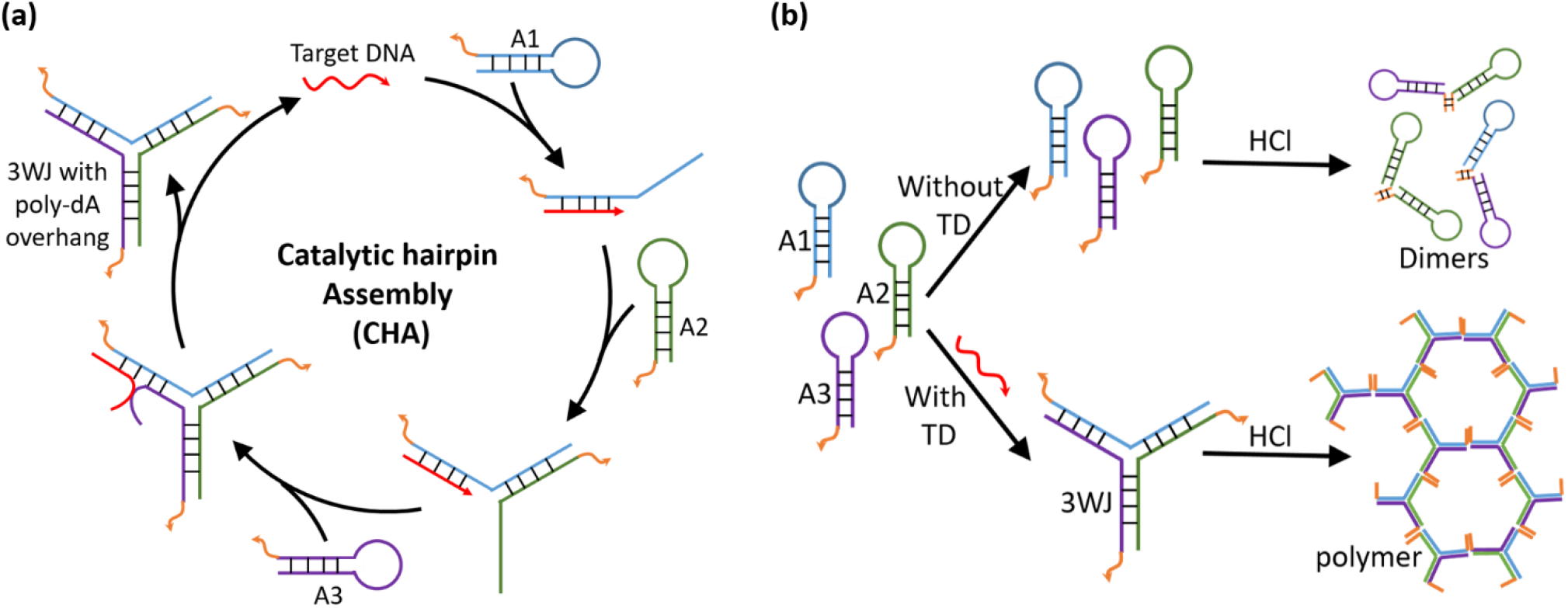
Schematic representation of (a) catalytic hairpin assembly driven formation of 3WJ and (b) polymer formation in presence of target DNA (TD).

The opening of hairpin-loop in presence of TD was analysed by EMSA, where the observed mobility was less in presence of TD (TD > A1 > A1+TD > A1+A2+TD > A1+A2+A3+TD) which confirms the 3WJ formation (**Figure 4a**). In the absence of TD, there is no opening of hairpin loop and strands A1, A2 and A3 remains as separate hairpin structures (Lane 7 in **Figure 4a**). The CHA leads to the formation of 3WJ and in presence of acid it forms polymer while in absence of TD, the strands remain as hairpin-loop form dimers only (**Scheme 1b**). The polymer formation in the presence of TD results in significant decrease (**Figure 4c**) in the liquid rise in the capillary tube in comparison to in absence of TD which can be seen with naked eyes (**Figure 4b**). The system is fully configurable, by changing the sequence in the monomer 3WJ, it can be re-designed against any pathogenic nucleic acid.

**Figure 4.**
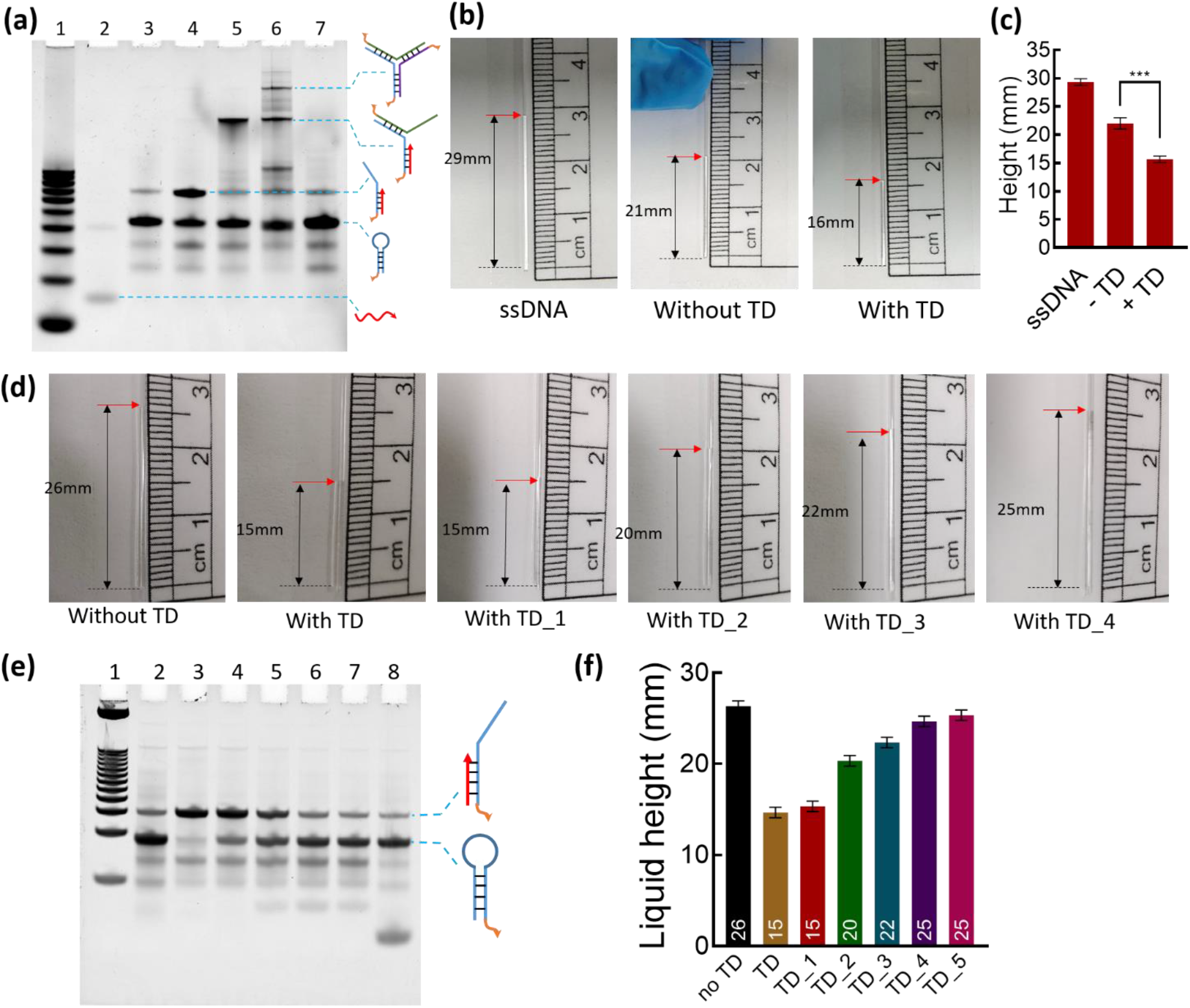
Visual detection of target DNA by capillary assay. (a) Gel-based characterization of formation of 3WJ by strand displacement; Lane 1=10bp DNA ladder, 2= TD, 3=A1, 4=A1+TD, 5=A1+A2+TD, 6=A1+A2+A3+TD, 7=A1+A2+A3. (b) Liquid height in the capillary tube at acidic pH with only one strand (A1) = 29 mm, all three strands (A1+A2+A3) = 21 mm and all three strands with target DNA (A1+A2+A3+TD) = 16 mm. (c) The comparison and quantification of liquid height from figure 4b. (d) Specificity of the assay towards the target DNA. (e) Affinity of A1 hairpin strand towards TD and its mutated versions analysed by EMSA; Lane 1=25bp DNA ladder, 2=A1, 3=A1+TD, 4=A1+TD_1, 5=A1+TD_2, 6=A1+TD_3, 7=A1+TD_4, 8=A1+TD_5. (f) The comparison and quantification of liquid height from figure 4d. Error bars indicate standard deviation; n = 3 (ordinary one-way ANOVA, P value < 0.001***).

#### 2.4.2 Specificity towards the target sequence

The capillary-based system was tested for specific detection of the TD for which it was designed. We used the different mutants of the TD, and analysed the liquid rise inside the capillary. We include the mutants with one (TD_1), two (TD_2), three (TD_3), four (TD_4) and complete base pair mismatch (TD_5) for the study. The binding efficiency of these mutants with A1 have been tested by EMSA, and found the decrease in the duplex formation with increase in the base pair mismatch (**Figure 4d**). That shows the capability of TD to initiate the CHA. In the capillary assay, the observed liquid rise in absence of any target DNA was 26 mm, while in presence of TD, TD_1, TD_2, TD_3, TD_4, and TD_5 it was 15 mm, 15mm, 20 mm, 22 mm, 25 mm, and 26 mm respectively (**Figure 4e & 4f**). This stipulate the sequence specificity of the designed system is up to two mismatched base pair. In other words, it can detect the difference of two or more nucleotides in the target sequence accurately (**Figure 5**).

**Figure 5.**
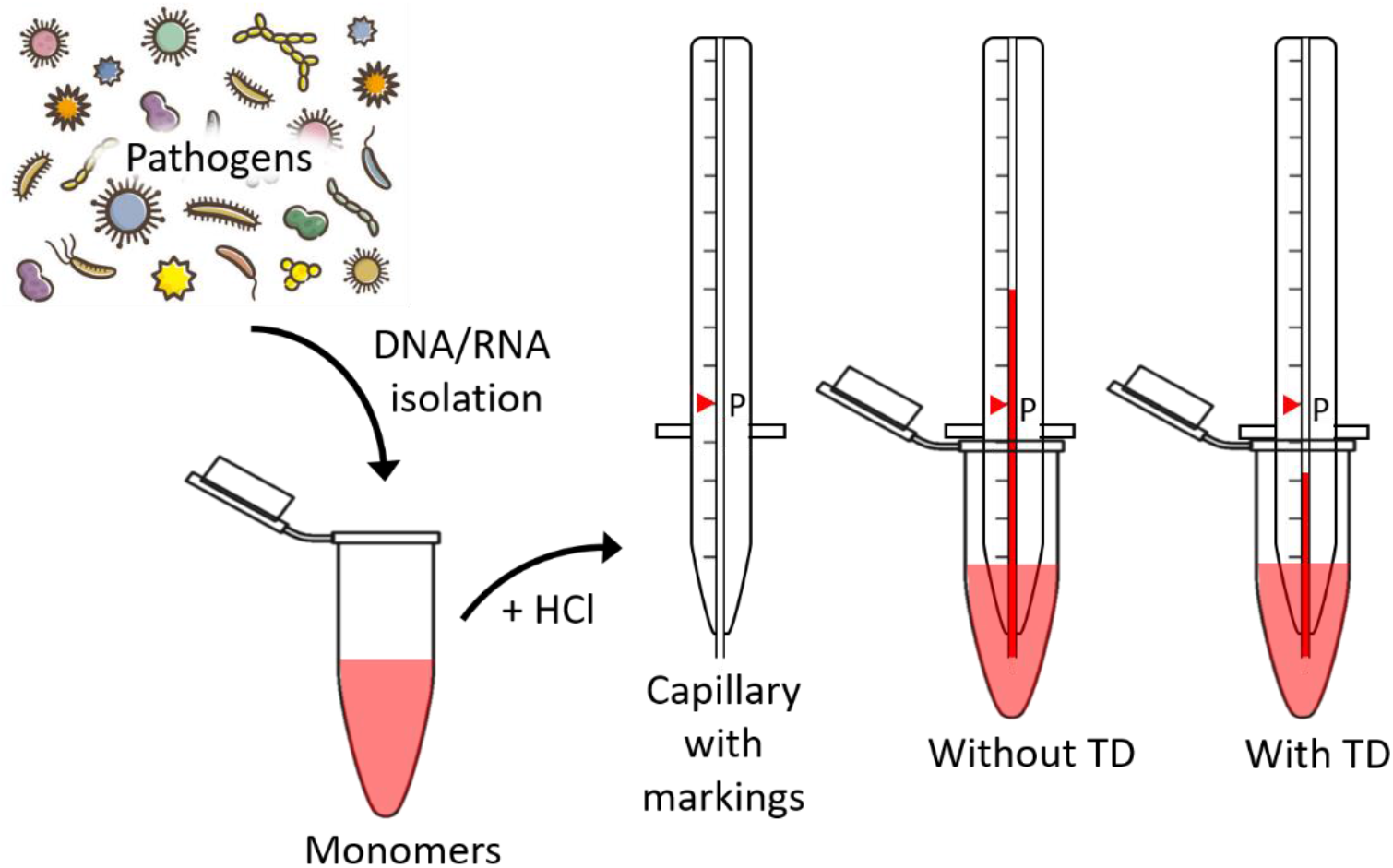
Schematic representation of the proposed capillary-action based visual detection system for pathogenic nucleic acid. The isolated pathogenic nucleic acid can be identified as targeted DNA, when introduced to poly-dA-3WJ monomers and then acidified the sample before dipping the marked capillary tube. The capillary is marked with threshold height, if liquid rises beyond this point the sample would be considered as negative and if it remains below point (formed polymer) then it would be positive.

## 3. Conclusions and discussion

For the first time, we have successfully utilized the pH-responsive A-motif to polymerize DNA 3WJ into supramolecular hydrogel. The formation of higher order structures at different pH was characterized by EMSA and DLS. At acidic pH, the retardation in mobility was seen in EMSA and larger structures in terms of hydrodynamic diameter was observed in DLS. The highly branched and porous morphology was imaged by SEM and AFM. Then the characteristic physical properties of a hydrogel were analysed by a rheometer, where sol-to-gel transformation can be seen as the change in the elastic (G′) and viscous response (G″). All the characterizations done on the A-motif hydrogel suggest that a pH dependent confirmation change is there.

The exclusive advantage of using A-motif as a cross-liker over other DNA motifs is there is no need for sequence design. Unlike other DNA motifs^25^, only a poly-dA chain of minimum six base pair is required to make an A-motif. We also inspect the *in-situ* formation of A-motif hydrogel, where the increase in the fluorescence intensity at acidic pH indicates the formation of hydrogel matrix on the cell surface. To explore further, we then employed the A-motif hydrogel in to a capillary based visual nucleic acid detection system (**Figure 5**), by considering the pH based physical changes from monomer to polymer. The designed detection system is sensitive up to single base pair mismatch. Also, it’s completely adaptable to different target nucleic acids, by changing the 3WJ forming sequence with complementary sequence of the desired target sequence. It was just to establish a proof-of-concept, to show the potential of the A-motif cross-linker. The current study demonstrates a new DNA motif which can be used further as a cross-linker or a functional nano-device. The A-motif have the ability for more real-world applications including controlled drug release, biosensing and many more to explore.

## 4. Materials and Methods

### Materials

All oligonucleotides (**Table S1**) at a 0.2 μM synthesis scale with desalting purification (HPLC purification in case of labelled ones) were purchased from Merck (Sigma-Aldrich). Mowiol® 4-88, Hoechst, Triton-X and loading dye were also purchased from Merck (Sigma-Aldrich). Acrylamide/Bis-acrylamide (29:1), Ethidium bromide, tetramethylethylenediamine (TEMED), paraformaldehyde and ammonium persulfate, Tris-acetate-EDTA (TAE) were purchased from HiMEdia. The cell culture dishes for adherent cells (treated surface) were procured from Nunc (Thermo Scientific). Dulbecco’s modified Eagle’s medium (DMEM), fetal bovine serum (FBS), penicillin−streptomycin and trypsin−ethylenediaminetetraacetic acid (EDTA) (0.25%) were purchased from Gibco (Thermo Scientific). Other salts and acid like MgCl_2_, NaCL, KCl, HCl, NaOH, CH_3_COONa, CH_3_COOH, Na_2_HPO_4_, and KH_2_PO_4_ were purchased from Merck.

### Synthesis of A-motif DNA hydrogel

Monomeric 3WJ were assembled from three single stranded DNA (ssDNA) via complementary base pairing. The mixture consisting of 100 μM of each stand i.e., dA-Y1, dA-Y2 and dA-Y3 with 2 mM MgCl_2_ were heated up to 95 °C and gradually cooled down till 20°C. The reaction cycle has a 5°C step decrease with an interval of 15 min at every step. The resulting mixture were having 3WJ monomers with poly-dA overhangs and remains like this at neutral pH. To form the hydrogel, 0.01 mM HCl (working concentration) were added to the monomer’s solution, so the final pH of the solution was 5 (pH = -log [H^+^]).

### Characterization

#### Electrophoretic Mobility Shift Assay

For EMSA, 10% native polyacrylamide gel electrophoresis (PAGE) was used. The samples for loading were diluted up to 2 μM with loading buffer (1X TAE), and mixed with 1X loading dye and kept for 3 min to allow the dye integration with the DNA completely. The samples were then loaded into the wells and was run at 10 volts/cm for 80 min in 1× TAE running buffer. For the different pH conditions i.e. pH 5 and pH 7, 1x (0.04 M) Britton–Robinson buffer (BRB) was used, in two different gel apparatus simultaneously. Finally, the gels were stained with ethidium bromide, and then scanned using a Bio-Rad ChemiDoc MP imaging system.

### Dynamic Light Scattering

The structural change from monomer (neutral pH) to polymer (acidic pH) and the reversible properties were analysed by change in the hydrodynamic diameter of the structures. A quartz cuvette of 50 μL working volume is used with Malvern Panalytical Zetasizer Nano ZS instrument. All the samples were centrifuged at 5,000 x g for 5 minutes to spin down any dust particles before analysis.

### Scanning electron microscope

The morphology of the hydrogel was characterized using a Jeol JSM-7600F field emission scanning electron microscope (FE-SEM) at an acceleration voltage of 5 kV. The samples were prepared by freeze-drying on to a glass substrate. Prior to imaging, platinum was coated on the sample via sputtering for 90 seconds to make the surface conductive.

### Atomic Force Microscopy Studies

The morphological characterization of synthesized A-motif DNA hydrogels was done on a Bruker Nano wizard Sense atomic force microscope (AFM) in tapping mode to collect the height data. AFM scanning was done via the peak force tapping mode to analyse the surface thickness, height, and roughness. A Si-doped antimony tip of 8 nm radius with spring constant of 3N/m and frequency 70KHz was used.

### Confocal Microscopy

The confocal laser scanning microscopy was done on a Leica TCS SP8 microscope. Cells were imaged after fixation using 63x oil immersion objective lens and the pinhole was set at 1 airy unit. Three lasers were used to excite the fluorophores i.e., 405 nm for Hoechst, 488 nm for GFP, and 633 nm for Cy5. The post processing was done on Fiji ImageJ 1.52p software^26,27^. For each condition 30–40 cells were quantified to study the in-situ formation of hydrogels.

### Rheological studies

The Rheology measurements were done on Anton Paar MCR702 rheometer. The monomer solution (neutral pH) was placed between parallel plates of the rheometer at 25°C and angular frequency range 0.1-100 rad s^−1^. The analysis was repeated with acidic condition at the same parameters.

### Cell culture

HeLa cells were cultured in DMEM supplemented with 10% FBS and 100 IU/mL penicillin−streptomycin. For all the studies at different pH, PBS (pH 7.4) and acetate buffer (pH 5) were used.

### Capillary assay

The assay is based on the capillary action of a capillary tube which is administered by the density of the liquid^28^. It is evident that a thin liquid will rise more in a capillary against gravity while a viscous liquid will not rise that much^29^. In case of A-motif hydrogel, the acidic pH would induce gelation and increase the viscosity of the liquid and lead to decrease in the liquid rise inside the capillary (**Figure S2**). We used glass capillary tubes with 0.5 mm inner diameter and 100 mm length for the study, and dip in to 100 μL of sample.

## Author’s contributions

DB and VM conceived the idea and planned the experiments. VM designed the experiments, performed most of the experiments and analyzed the data, wrote the first draft of the manuscript. AS helped with the rheology measurements. VM, CG, DB analyzed the data and helped with final draft of the manuscript.

## Acknowledgment

We sincerely thank the members of DB and CG labs for constant support and discussions. VM and AS thank IITGN, MoE GoI for the PhD fellowships. This work was supported by Gujarat State Biotechnology Mission, GSBTM. DB thanks SERB-DST GoI for Ramanujan Fellowship. The central instrumentation facilities at IITGN are gratefully acknowledged.

