## Supplementary information for "pH - responsive, reversible A-motif based DNA hydrogels: synthesis and biosensing applications"

**Table S1**: DNA oligos used for the study.

| Oligo Name | Sequence (5' -> 3') |
|---|---|
| dA Y1 | AAA AAA AAA AAA CGA CCG ATG AAT AGC GGT CAG ATC CGT ACC TAC TCG |
| dA Y2 | AAA AAA AAA AAA CGA GTC GTT CGC AAT ACG ACC GCT ATT CAT CGG TCG |
| dA Y3 | AAA AAA AAA AAA CGA GTA GGT ACG GAT CTG CGT ATT GCG AAC GAC TCG |
| dA Y1-Cy5 | Cy5-AAA AAA AAA AAA CGA CCG ATG AAT AGC GGT CAG ATC CGT ACC TAC TCG |
| A1 | AAA AAA AAA AAA T GGA ACA ACA TTG CCA GTC TCG ACT GCA TGG CAA TGT TGT TCC TTG AGG AAG |
| A2 | AAA AAA AAA AAA T CCA GTC TCG ACT GCA CTT CCT CAA GGA TGC AGT CGA GAC TGG CAA TGT TGT |
| A3 | AAA AAA AAA AAA T GCA CTT CCT CAA GGA ACA ACA TTG CCA TCC TTG AGG AAG TGC AGT CGA GAC |
| TD | CTT CCT CAA GGA ACA ACA TTG CCA |
| TD_1 | CTA CCT CAA GGA ACA ACA TTG CCA |
| TD_2 | CTA CCT CAT GGA ACA ACA TTG CCA |
| TD_3 | CTA CCT CAT GGA ACT ACA TTG CCA |
| TD_4 | CTA CCT CAT GGA ACT ACA CTG CCA |
| TD_5 | GCA TGG CAC TCC TTG TGT ACA TGT |

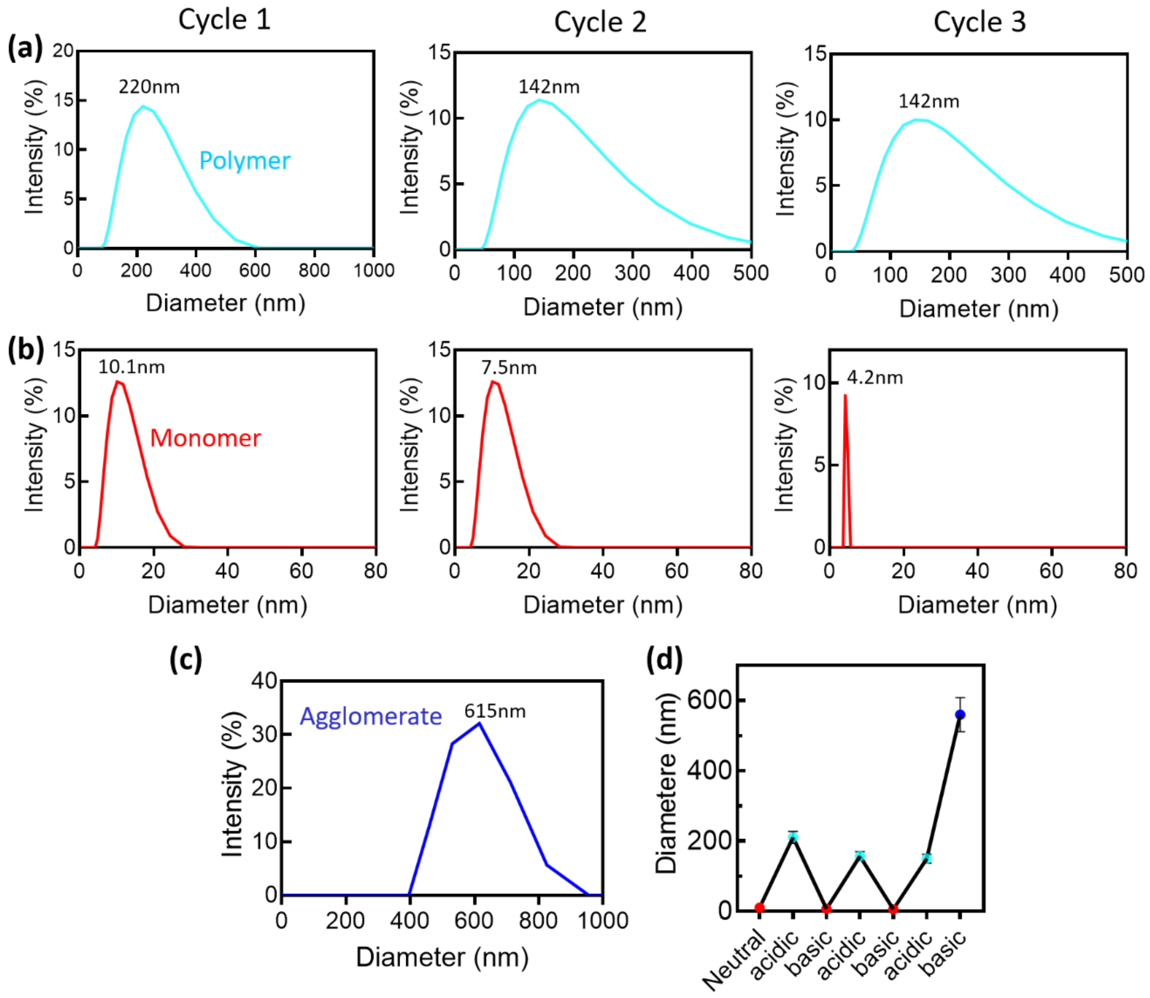


**Figure S1**. Reversibility of A-motif hydrogel. (a) hydrodynamic diameters (>100 nm) at acidic conditions in three alternate acid-base cycles. (b) hydrodynamic diameters (<10 nm) at basic conditions in three alternate acid-base cycles. (c) Aggregate formation after the third cycle. (d) Comparison of hydrodynamic diameters at different pH cycles.

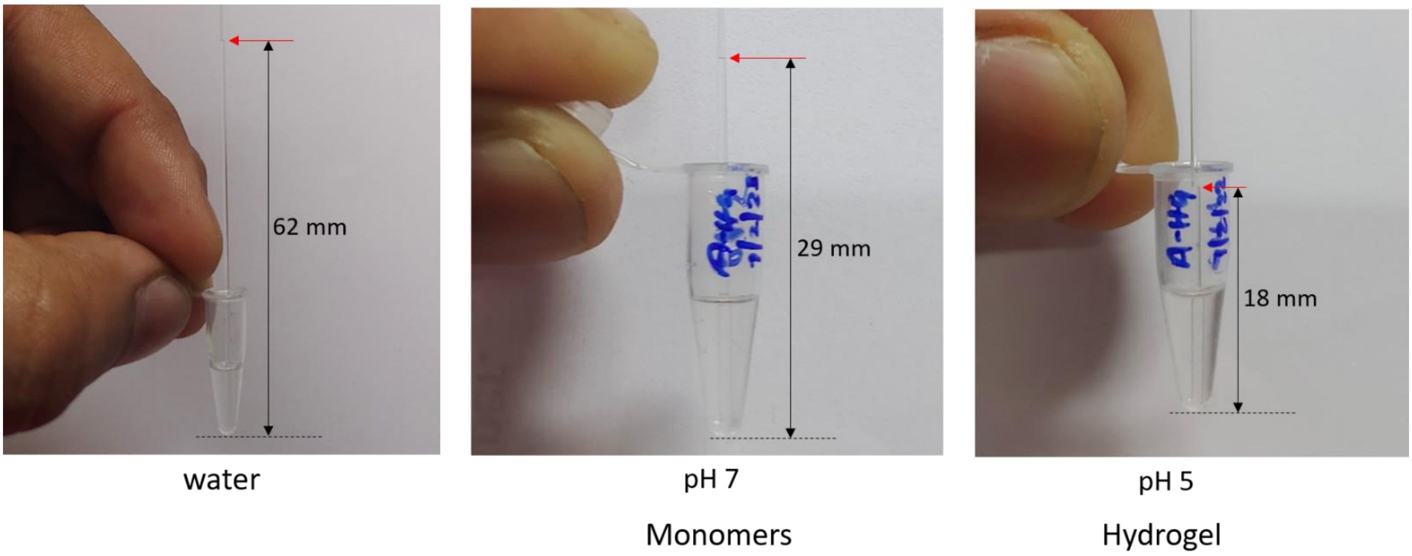


**Figure S2**. The visuals of pH-dependent capillary action.
